# Sortilin inhibition treats multiple neurodegenerative lysosomal storage disorders

**DOI:** 10.1101/2023.09.22.559064

**Authors:** Hannah G. Leppert, Joelle T. Anderson, Kaylie J. Timm, Cristina Davoli, Melissa A. Pratt, Clarissa D. Booth, Katherine A. White, Mitchell J. Rechtzigel, Brandon L. Meyerink, Tyler B. Johnson, Jon J. Brudvig, Jill M. Weimer

## Abstract

Lysosomal storage disorders (LSDs) are a genetically and clinically diverse group of diseases characterized by lysosomal dysfunction. Batten disease is a family of severe LSDs primarily impacting the central nervous system. Here we show that AF38469, a small molecule inhibitor of sortilin, improves lysosomal and glial pathology across multiple LSD models. Live-cell imaging and comparative transcriptomics demonstrates that the transcription factor EB (TFEB), an upstream regulator of lysosomal biogenesis, is activated upon treatment with AF38469. Utilizing CLN2 and CLN3 Batten disease mouse models, we performed a short-term efficacy study and show that treatment with AF38469 prevents the accumulation of lysosomal storage material and the development of neuroinflammation, key disease associated pathologies. Tremor phenotypes, an early behavioral phenotype in the CLN2 disease model, were also completely rescued. These findings reveal sortilin inhibition as a novel and highly efficacious therapeutic modality for the treatment of multiple forms of Batten disease.

## Introduction

Lysosomal storage disorders (LSDs) are a clinically and genetically diverse group of diseases characterized by lysosomal dysfunction. Pathogenic variants in more than 70 genes result in LSDs characterized by disruptions in lysosomal biogenesis, homeostasis, or catabolic function.^1^ LSDs vary widely in terms of the affected tissues, timing of onset and progression, and clinical severity. Many are fatal and severely impact quality of life. While each LSD is rare (ranging from 1 in 50,000 to 1 in 250,000 live births, or fewer), LSDs are collectively quite common, with an overall incidence of at least 1 in 5,000 live births^1, 2^. Due in part to their individual rarity and etiological heterogeneity, most LSDs lack an FDA-approved disease modifying treatment, leaving patients with substantial unmet need. One subset of LSDs, Batten disease (also referred to as neuronal ceroid lipofuscinosis), are a family of diseases characterized by lysosomal dysfunction leading to the accumulation of ceroid lipofuscin (autofluorescent storage material, ASM) and cellular dysfunction and degeneration throughout the central nervous system.^3^

The phenotypically similar Batten disease variants are predominantly pediatric and rare with an incidence of 2–4/100,000 births and are most commonly fatal.^4, 5^ Primarily autosomal recessive, Batten disease is caused by mutations in one of at least 13 known **<u>c</u>**eroid **l**ipofuscinosis, **<u>n</u>**euronal (*CLN*) genes that encode for a variety of extralysosomal and lysosomal proteins, many of which have unclear functions.^6^ Despite the variability in causative genetics, etiology, and symptomatology, all forms of Batten disease cause neurodegenerative CNS-related symptoms such as progressive mental and motor deficits, vision loss and seizures. Several disease-modifying gene therapies for CLN2, CLN3, CLN5, CLN6, and CLN7 diseases have been introduced into clinical trials, some with promising results.^7–14^ However, the limited transduction (10-50% of cells) achieved with contemporary viral vectors will limit efficacy for these cell-autonomous disorders. Furthermore, microglia, a key brain cell type for Batten disease pathogenesis, are entirely refractory to AAV transduction and thus remain unaddressed with most existing gene therapies.^15–17^ Only one form of Batten disease, CLN2 disease, has an FDA-approved disease-modifying treatment – an enzyme replacement therapy called cerliponase alfa (Brineura®), which is well-tolerated and stabilizes language and motor abilities but fails to halt gray matter loss in the brain.^18, 19^ While gene therapy and enzyme replacement therapy offer encouraging results, limitations such as incomplete transduction, patient immune-responses, and bioavailability emphasize an overwhelming need for more effective therapies.^20, 21^ ^4^

Small molecule therapeutics have been shown to have some efficacy in LSDs such as Fabry disease, Gaucher disease, and Pompe disease as well as neurogenerative diseases such as Alzheimer’s, Parkinson’s, and Huntington’s disease.^22–29^ For Batten disease, an effective small-molecule therapy must address a complex etiological milieu including primary defects in lysosomal function and downstream neuroinflammatory and neurodegenerative cascades. One promising strategy for restoration of lysosomal function is small-molecule-mediated activation of transcription factor EB (TFEB), a master regulator of lysosomal biogenesis. ^30–34^ TFEB activates the Coordinated Lysosomal Expression and Regulation (CLEAR) network, a family of lysosomal and autophagy-related genes containing CLEAR motifs in regulatory regions. TFEB activation has been shown to improve LSD pathology through improving cellular clearance via lysosomal biogenesis, autophagosome formation, and lysosomal-autophagosome fusion.^31, 35, 36^ Small molecule TFEB activators have been shown to have some efficacy *in vivo* in models of Batten disease due to enhancement of cellular clearance, but neuroinflammation remains largely unaddressed with such a strategy.^30, 31^ Conversely, small molecule therapies aimed at reducing neuroinflammation have been shown to have some benefit in Batten disease mouse models, but lysosomal dysfunction remains unaddressed.^37^

Here, we demonstrate that small molecule antagonism of sortilin (SORT1) enhances lysosomal function through TFEB activation while simultaneously ameliorating neuroinflammation with direct effects on microglia. Sortilin, a member of the vacuolar protein sorting 10 protein (Vps10p) domain receptor family, is a systemic, multifunctional protein that has been implicated in the etiology of Alzheimer’s disease, diabetes, cardiovascular disease, frontotemporal dementia, auto-inflammatory and pro-inflammatory disorders as well as various cancers.^38–43^ Sortilin was first described as an intracellular sorting protein, directing cargoes to various endocytic compartments including the lysosome.^44–46^ Sortilin is widely distributed throughout all neuronal cell populations. At the cellular level, sortilin is primarily localized to the trans-Golgi network (TGN) and other intracellular membrane bound organelles and is involved in lipid metabolism, lysosomal protein sorting, and neurotrophic signaling.^47–51^ Using a high-content screening pipeline with cell models of five etiologically diverse forms of Batten disease (CLN1, CLN2, CLN3, CLN6, CLN8), we identified the SORT1 antagonist AF38469 as a potent molecule for reducing lysosomal storage material. We subsequently demonstrate the efficacy of sortilin inhibition in several *in vitro* and *in vivo* mouse models of Batten disease, showing that treatment with the small molecule AF38469 reduces lysosomal dysfunction, increases the production of lysosomal enzymes, and operates at least partially through TFEB and downregulation of inflammatory cytokines.

## Results

### Sortilin inhibition potently reduces lysosomal storage material accumulation in multiple cell models of lysosomal storage disorders

To determine if the small molecule sortilin inhibitor AF38469 was efficacious in cellular models of Batten disease, the compound was screened in cultured mouse embryonic fibroblasts (MEFs) and primary cortical neurons (PNCs) of five Batten disease mouse models. These models have diverse disease etiologies: *Cln1^R^*^151^*^X^* with a mutation in the lysosomal palmitoyl thioesterase PPT1^52^, *Cln2^R^*^207^*^X^* with a mutation in the lysosomal protease TPP1^53^, *Cln3^Δex^*^7^*^/^*^8^ with a mutation in the transmembrane endosomal/lysosomal protein CLN3^54^*, Cln6^nclf^* with a mutation in the transmembrane ER-resident CLN6 protein^55^, and *Cln8^mnd^* with a mutation in the transmembrane ER/Golgi protein CLN8^56^. While primary etiologies differ markedly across different forms of Batten disease, downstream cellular pathologies are remarkably consistent and are characterized by storage of numerous lysosomal substrates (including those not directly catabolized by the causative proteins), neuroinflammation, and neurodegeneration.

Accumulation of autofluorescent storage material (ASM) is a hallmark found in all forms of Batten disease.^57^ Treatment with AF38469 significantly reduced the accumulation of ASM in *Cln1^R^*^151^*^X^*, *Cln2^R^*^207^*^X^*, *Cln3^Δex^*^7^*^/^*^8^, *Cln6^nclf^*, and *Cln8^mnd^* MEFs (**Figure 1A-F, Supplemental Figure 1A-F**). Similarly, the accumulation of ASM was significantly reduced by treatment with AF38469 in *Cln1^R^*^151^*^X^*, *Cln2^R^*^207^*^X^*, *Cln3^Δex^*^7^*^/^*^8^, and *Cln8^mnd^* PNCs (**Figure 1A-D, F, Supplemental Figure 2A-F)**. However, ASM accumulation was significantly increased in *Cln6^nclf^*PNCs (**Figure 1E**). Since sortilin inhibition is known to increase the availability of the lysosomal enzyme chaperone progranulin (GRN), we asked whether AF38469 could still reduce ASM in the context of GRN deficiency. We treated GRN knockout (also referred to as *Cln11^-/-^*, given mutations in *GRN* have been classified as a variant of Batten disease) fibroblasts with the same doses of AF38469, but did not observe similar decreases in ASM, suggesting that sortilin inhibition reduces ASM in a GRN-dependent manner (**Figure 1G**).

**Figure 1.**
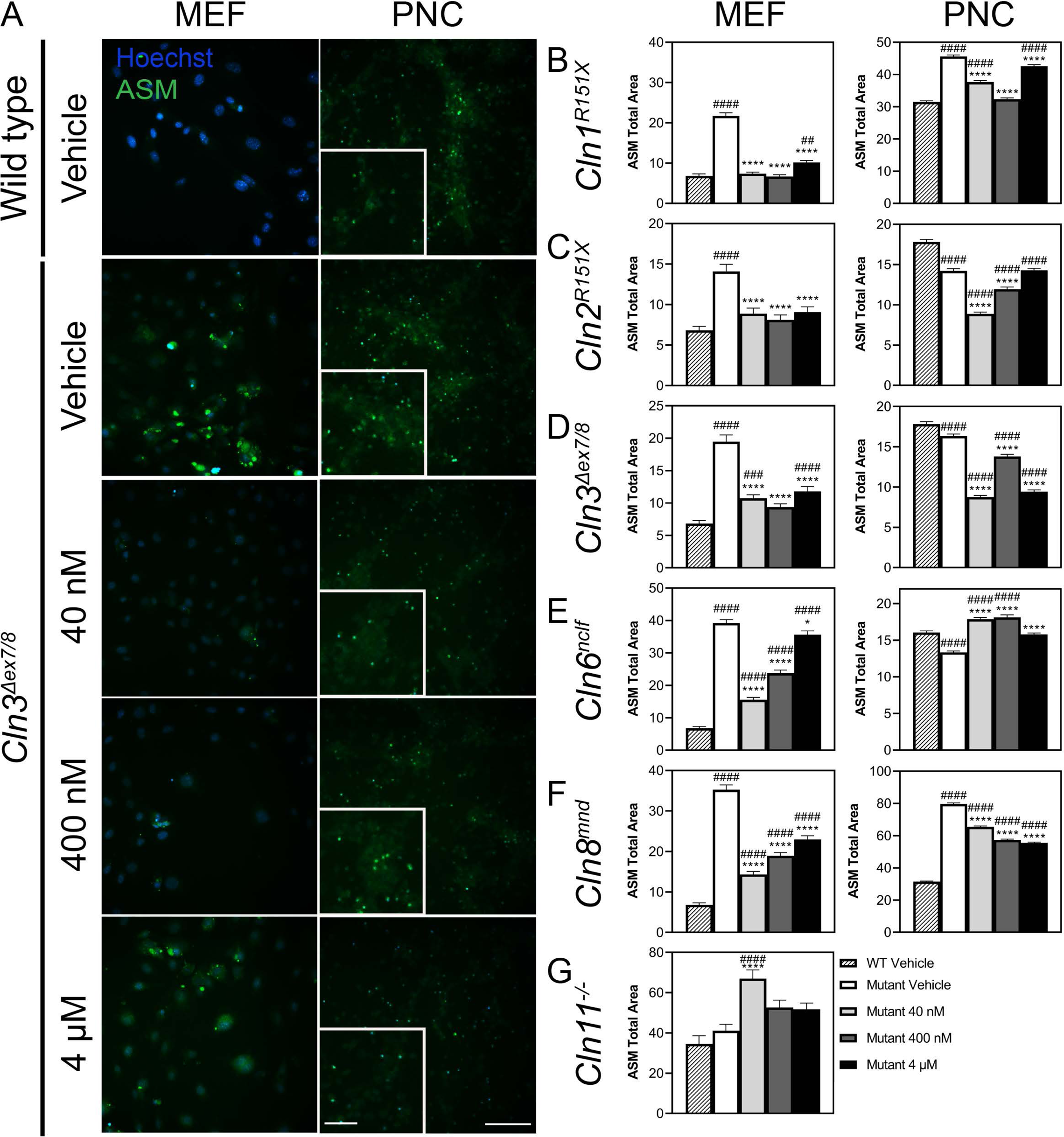
Inhibition of sortilin through AF38469 (40 nM, 400 nM, 4μM) impacts the accumulation of ASM in cell models of Batten disease. Mouse embryonic fibroblasts (MEFs) and primary cortical neurons (PNCs) were isolated from E15.5 embryos from *Cln1^R^*^151^*^X^*, *Cln2^R^*^207^*^X^*, *Cln3^Δex^*^7^*^/^*^8^*, Cln6^nclf^*, and *Cln8^mnd^*mouse lines. *Cln11^-/-^* fibroblasts were isolated from adult *Cln11^-/-^* mice. All cell lines were dosed with drug-containing media on day *in vitro* (DIV) 3 and DIV5 and analyzed on DIV7 using the CellInsight CX7 High-Content Screening Platform (CX7) (A) Representative ASM accumulation in *Cln3^Δex^*^7^*^/^*^8^ MEFs and PNCs. Upon treatment with AF38469 (40 nM, 400 nM, 4μM), ASM accumulation was (B-F) reduced in *Cln1^R^*^151^*^X^ Cln2^R^*^207^*^X^*, *Cln3^Δex^*^7^*^/^*^8^*, Cln6^nclf^*, and *Cln8^mnd^* MEFs; (B-D, F) reduced in *Cln1^R^*^151^*^X^ Cln2^R^*^207^*^X^*, *Cln3^Δex^*^7^*^/^*^8^, and *Cln8^mnd^* PNCs; and (E, G) increased in *Cln6^nclf^*PNCs and *Cln11^-/-^* MEFs. Mean ± S.E.M.. One-way ANOVA with a Šidák post-hoc test, *^/#^p<0.05, **^/##^p<0.01, ***^/###^p<0.001, ****^/####^p<0.0001. Hashsigns indicate comparison to WT, asterisks indicate comparison to mutant vehicle. MEFs and PNCs n=1000-4000 and 4000-20000 cells/treatment group, respectively. Scale bar, 100 μm, Inset scale bar, 50 μm.

In cell models of lysosomal storage disorders, accumulation of lysosomal storage material often results in the excessive accumulation of dysfunctional lysosomes within the cell. We also examined the effect of AF38469 on lysosomal accumulation using Lysotracker, a pH-dependent marker of lysosomes. While phenotypes were more variable, we observed a reduction in signal in most of our cell lines, supporting a net resolution of dysfunctional lysosomes (**Supplemental Figure 1H-N, Supplemental Figure 2G-L**). We observed no impact on cellular viability upon treatment (**Supplemental Figure 1O-U, Supplemental Figure 2M-R**). Interestingly, the observed ASM and lysotracker effects differed markedly with dose, with the lower 40 nM dose generally providing the most benefit, suggestive of a hormetic response.^58^ Taken together, these data indicate that sortilin inhibition through AF38469 reduces lysosomal pathology in a wide range of NCL cell models without impacting cellular viability.

### AF38469 stimulates TFEB/TFE3 nuclear translocation and transcription of CLEAR network genes

Since sortilin inhibition reduced the accumulation of lysosomal storage material resulting from diverse etiologies, we asked whether this could be due in part to the activation of transcriptional networks impacting lysosomal function – namely the coordinated lysosomal expression and regulation (CLEAR) network controlled by transcription factor EB (TFEB) and transcription factor binding to IGHM enhancer 3 (TFE3). To test this hypothesis, we cultured wild type Neuro 2A cells expressing plasmids encoding TFEB or TFE3 fused with GFP and monitored the nuclear translocation of the transcription factors with live imaging after being treated with AF38469. Increased TFEB and TFE3 nuclear translocation was observed within 90 minutes of a low-dose treatment of AF38469 (**Figure 2A**), suggesting that SORT1 inhibition may have initiated a hormetic stress-response wherein blockade of the lysosomal receptor was compensated for by upregulation of lysosomal biogenesis.

**Figure 2.**
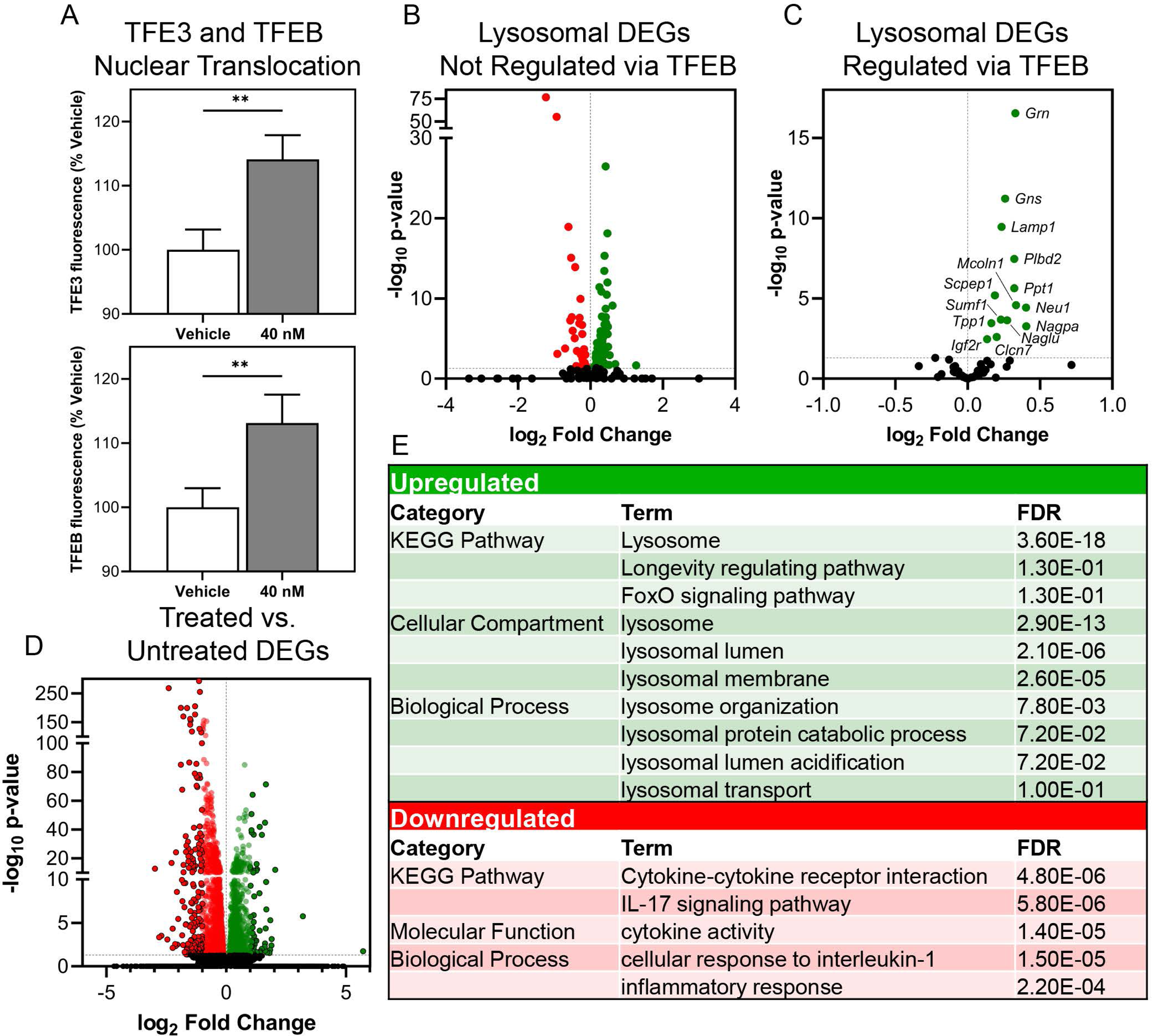
Inhibition of sortilin through acute treatment with AF38469 stimulates TFEB and TFE3 nuclear translocation and upregulation of TFEB gene targets. Wild type Neuro 2A cells were transfected with pEGFP-N1-TFEB or pEGFP-N1-TFE3 and acutely treated with vehicle or AF38469 (40 nM). (A) Treatment with AF38469 significantly stimulated TFEB and TFE3 nuclear translocation after 90 minutes of treatment as measured by live cell imaging. Expression analysis of lysosomal differentially expressed genes (DEGs) in wild type AF38469 (40 nM) treated MEFs compared to vehicle treated (B) showing up and downregulation of lysosomal DEGs not regulated by TFEB compared to (C) showing distinct pattern of upregulation in TFEB targeted lysosomal DEGs organized by adjusted p-value and fold change. (D) Expression analysis of all DEGs. (E) Table of select enriched upregulated and downregulated DAVID GO terms from DEG analysis. Mean ± S.E.M. One-way ANOVA, Dunnett’s multiple comparisons. *p <0.05, **p<0.01, ****p<0.0001. A: n=5000-6500 cells/treatment. For B-F: RNA collected from 3 wells/treatment. Adjusted p-value and fold change.

To confirm that downstream targets of TFEB and TFE3 were activated following treatment with AF38469, we treated wild type MEFs with DMSO or AF38469 (40 nM) and assessed the expression of differentially expressed lysosomal genes.^59, 60^ When analyzing all lysosomal genes and lysosomal genes not known to be regulated by TFEB, we saw no pattern of altered regulation in AF38469 treated wild type MEFs (**Figure 2B, C**). However, when analyzing TFEB target genes in isolation, 100% of the differentially expressed genes were upregulated by drug treatment (**Figure 2D**). The top hits in terms of fold change and significance encoded a range of lysosomal enzymes (e.g. *Ppt1*, *Tpp1*, *Naglu*, etc.), lysosomal trafficking proteins (e.g. *Igf2r*, *Mcoln1*) and transmembrane proteins critical for lysosomal homeostasis (e.g. *Lamp1*, *Clcn7*) (**Figure 2E,F**).^61–66^ Pathway analysis demonstrated enrichment in upregulated DEGs for lysosome compartments. Enrichment of downregulated DEGs revealed pathways involved in cytokine interactions and pathways as well as inflammation. Collectively, these data indicate that inhibition of SORT1 influences lysosomal gene regulation through the CLEAR network.

### Treatment with AF34869 increases TPP1 and PPT1 enzyme activities in Batten disease cell lines

In Batten disease, lysosomes have reduced catabolic function, and lysosomal proteins such as PPT1 (the protein encoded by *CLN1*) and TPP1 (the protein encoded by *CLN2*) have altered function.^67, 68^ Since AF38469 significantly upregulated the expression of both of these genes, we performed TPP1 and PPT1 enzyme activity assays on Batten disease MEFs treated with AF38469 (40 nM). Drug treatment significantly increased PPT1 enzyme activity levels in *Cln2^R^*^207^*^X^*, *Cln3^Δex^*^7^*^/^*^8^, *Cln6^nclf^,* and *Cln11^-/-^* MEFs, restoring *Cln3^Δex^*^7^*^/^*^8^ MEFs activity level to that of wild type vehicle treated cells (**Supplemental Figure 3A-D)**. Treatment with AF38469 (40 nM) significantly increased TPP1 enzyme activity of *Cln3^Δex^*^7^*^/^*^8^, *Cln6^nclf^,* and *Cln11^-/-^*MEFs (**Supplemental Figure 3G-I**). As expected, no significant difference was seen in *Cln2^R^*^207^*^X^* MEFs due to the absence of protein product made from the mutated *Cln2* gene (**Supplementary Figure 3F)**.^53^ We also asked whether treatment with AF38469 would impact lysosomal function in wild type cells and observed significant increases in TPP1 and PPT1 enzyme activity (**Supplemental Figure 3E, J**). Overall, inhibition of sortilin through AF38469 improved lysosomal function in a genotype-independent manner as measured by PPT1 and TPP1 activity levels.

### Short-term treatment with AF38469 improves histopathological and behavioral outcomes in a CLN2 disease mouse model

We next asked whether the benefits of sortilin inhibition could translate *in vivo* and reduce histopathological and behavioral markers of disease severity in a mouse model of a particularly severe, infantile-onset LSD, CLN2 disease. The *Cln2^R^*^207^*^X^* mouse model recapitulates a common human pathogenic variant in *TPP1* leading to the absence of TPP1 enzyme activity (a lysosomal protease) and subsequent lysosomal storage and dysfunction. *Cln2^R^*^207^*^X^* mice exhibit severe, early-onset brain pathology, tremors, and reduced lifespan with most animals succumbing to the disease very rapidly, by 4 months of age, allowing us to screen for behavioral impacts in a short-term efficacy study.^53^

Homozygous *Cln2^R^*^207^*^X^* and littermate wild type mice received continuous treatment with AF38469 or vehicle via drinking water starting at wean until 11 weeks of age. Using doses found in previous AF38469 studies for unrelated disorders, we initially treated the mice with 3 μg/ml and 78 μg/ml doses which translated to roughly 10 μg and 250 μg per day.^69^ While both doses provided a complete rescue of tremor phenotypes in moribund animals (data not shown), we observed no benefit on histopathological outcomes (**Supplementary Figure 4A**). Since sortilin has important roles for lysosomal biogenesis, we hypothesized that the high-level inhibition provided by the 3 μg/ml and 78 μg/ml doses could be suboptimal. We thus enrolled an additional study with lower doses of 0.03 μg/ml and 0.3 μg/ml (0.1 μg, 1 μg per day, respectively) (**Figure 3A**).

**Figure 3.**
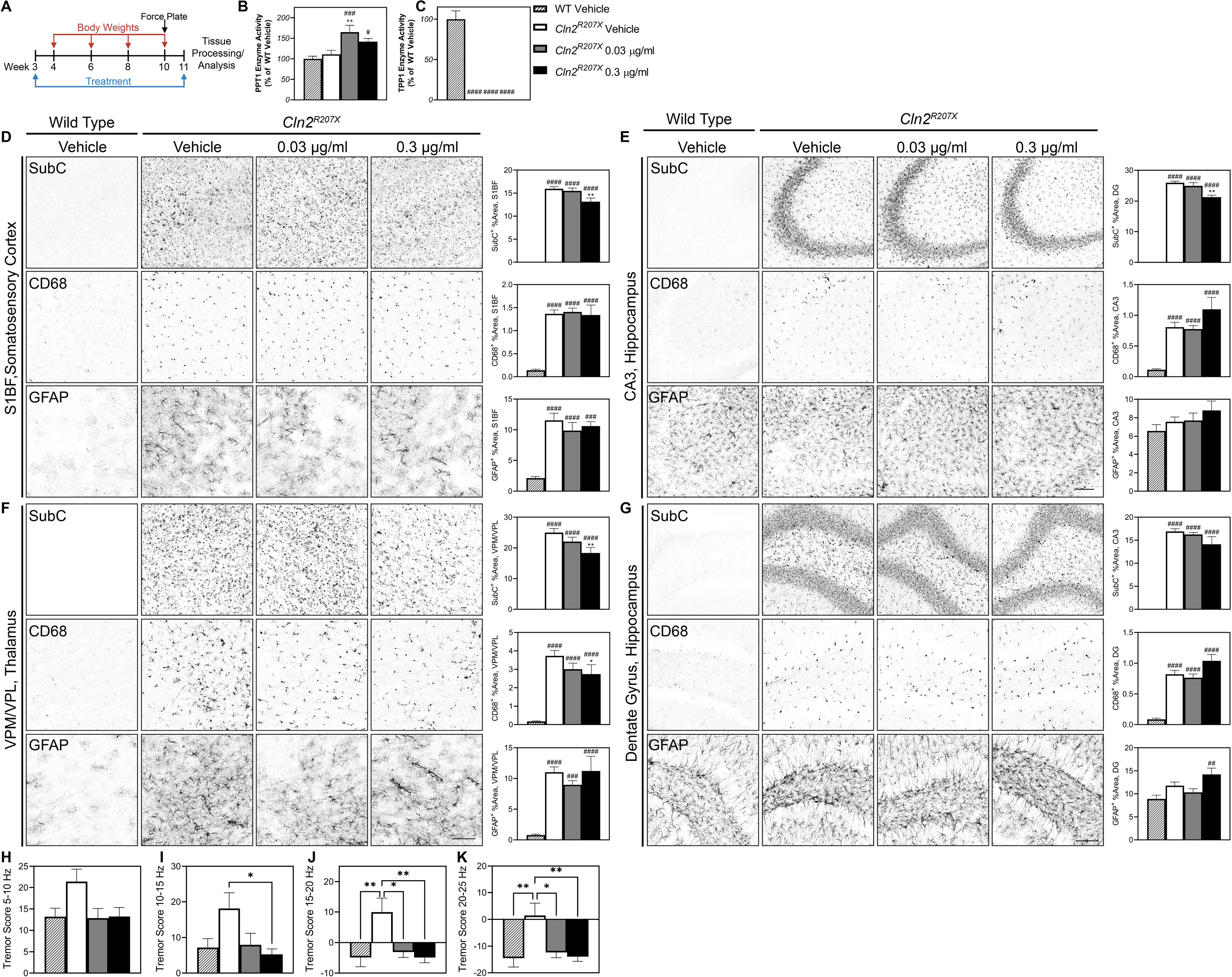
Short-term treatment with AF38469 improves histopathological and behavioral outcomes in *Cln2^R^*^207^*^X^* mice. (A) Homozygous *Cln2^R^*^207^*^X^* and littermatewild type mice received continuous treatment with AF38469 or vehicle via drinking water starting at wean until 11 weeks of age. (B) Treatment with AF38469 (0.03 μg/ml) significantly increased PPT1 enzyme activity levels in *Cln2^R^*^207^*^X^* treated mice. (C) Treatment with AF38469 had no impact on TPP1 enzyme activity levels *Cln2^R^*^207^*^X^* treated mice. (D, E) AF38469 treatment (0.3 μg/ml) in *Cln2^R^*^207^*^X^* mice significantly prevented SubC accumulation and had no impact on microglial reactivity (CD68^+^) or astroglial activation (GFAP^+^) in the S1BF of the somatosensory cortex and CA3 of the hippocampus. (F) Treated *Cln2^R^*^207^*^X^* mice (0.3 μg/ml) saw significant prevention of accumulation of mitochondrial ATP synthase subunit C (SubC^+^) and reduction of microgliosis (CD68^+^) in the VPM/VPL of the thalamus with no impact on astrocytosis (GFAP^+^). (G) Treatment with AF38469 had no impact on CLN2 Batten disease pathology in the dentate gyrus and hilus regions of the hippocampus. (H-K) Tremor index scores were reduced and restored to wild type levels in AF38469 treated *Cln2^R^*^207^*^X^*mice when compared to vehicle treated mice. Mean ± S.E.M. (B-G) Nested one-way ANOVA with a Šidák post-hoc test (H-K) One-way ANOVA with a Šidák post-hoc test *^/#^p<0.05, **^/##^p<0.01, ***^/###^p<0.001, ****^/####^p<0.0001. Hashsigns indicate comparison to WT, asterisks indicate comparison to mutant vehicle. Scale bar, 100 μm. n = 6-8 animals/treatment.

Neuropathology in various forms of Batten disease can be monitored through histopathological markers of lysosomal storage burden and reactive gliosis.^70–73^ To determine whether sortilin inhibition enhanced lysosomal function in the AF38469 treated *Cln2^R^*^207^*^X^* mice, we measured PPT1 and TPP1 enzyme activity in brain lysates. Treatment with AF38469 increased PPT1 enzyme activity and did not impact TPP1 activity as expected in the CLN2 mouse model (**Figure 3B,C**).^53^ We examined hallmark neuroinflammation in brain regions known to be involved in Batten pathogenesis, namely the ventral posterolateral and ventral posteromedial nuclei (VPM/VPL) of the thalamus, the somatosensory cortex barrel field (S1BF), the CA3 region of the hippocampus, and the dentate gyrus (DG) and hilus regions of the hippocampus.^74–76^ In *Cln2^R^*^207^*^X^* mice, treatment with AF38469 (0.3 μg/ml) slightly reduced levels of mitochondrial ATP synthase subunit C (SubC), a primary constituent of the lysosomal storage material, in the S1BF of the somatosensory cortex, CA3 of the hippocampus, and VPM/VPL of the thalamus (**Figure 3D-F**).^77^ *Cln2^R^*^207^*^X^* mice treated with AF38469 had reduced levels of microglial reactivity (CD68^+^) in the VPM/VPL of the thalamus while no impact was observed in other regions (**Figure 3F**). We observed no impact on reduction astroglial activation (GFAP^+^) in any region of the brain ^78, 79^ (**Figure 3D-G**). Collectively, histopathology demonstrated a reduction in lysosomal pathology and downstream neuroinflammation, suggesting an overall reduction of disease burden in the brains of treated animals.

To investigate whether these pathological benefits improved clinical symptoms, we examined seizure index scores using a force plate actimeter.^53^ AF38469 treatment at both 0.03 μg/ml and 0.3 μg/ml rescued *Cln2^R^*^207^*^X^*tremor index scores to wild type levels at frequency bands encompassing 5-10 Hz, 10-15 Hz, 15-20 Hz, and 20-25 Hz (**Figure 4H-K**). We additionally monitored body weights and determined that there was no toxic effect due to treatment with AF38469 (**Supplemental Figure 7A**).

**Figure 4.**
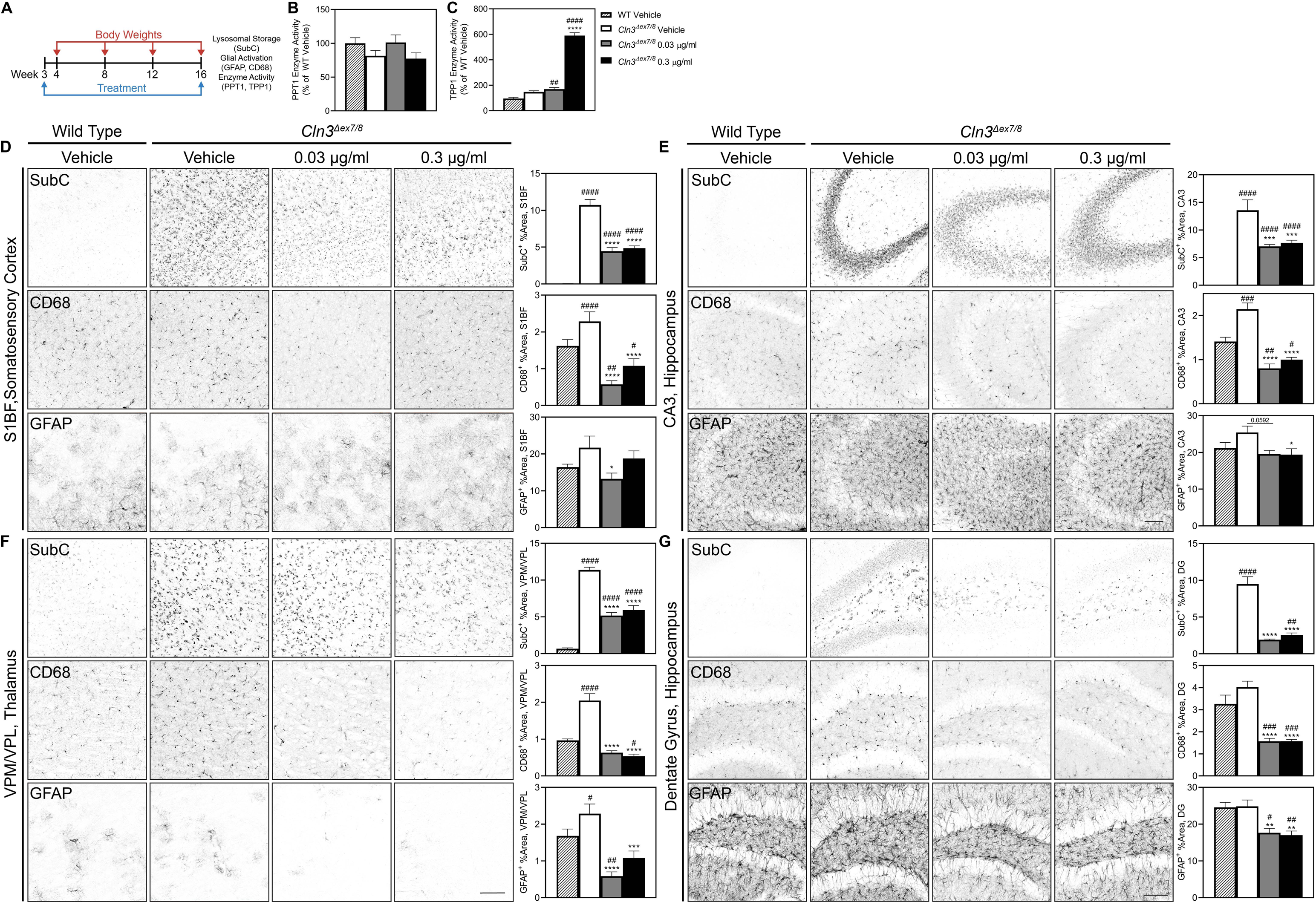
Short-term treatment with AF38469 improves histopathological outcomes in *Cln3^Δex^*^7^*^/^*^8^ mice. (A) Homozygous *Cln3^Δex^*^7^*^/^*^8^ and wild type mice received continuous treatment with AF38469 or vehicle via drinking water starting at wean until 16 weeks of age. B) Treatment with AF38469 had no impact on PPT1 enzyme activity levels in *Cln3^Δex^*^7^*^/^*^8^ treated mice (C) Treatment with AF38469 significantly increased TPP1 enzyme activity levels in *Cln3^Δex^*^7^*^/^*^8^ treated mice. (D, E) In the S1BF of the somatosensory cortex and CA3 of the hippocampus, AF38469 treatment in *Cln3^Δex^*^7^*^/^*^8^ mice significantly prevented SubC accumulation and microgliosis (CD68^+^). Treatment with the low and high dose significantly prevented astrocyte activation in the S1BF and CA3, respectively. (F, G) Treated *Cln3^Δex^*^7^*^/^*^8^ mice saw significant prevention of accumulation of mitochondrial ATP synthase subunit C (SubC^+^) and reduction of microgliosis and astrocytosis (CD68^+^, GFAP^+^) in the VPM/VPL of the thalamus and dentate gyrus and hilus regions of the hippocampus. Mean ± S.E.M. Nested one-way ANOVA with a Šidák post-hoc test *^/#^p<0.05, **^/##^p<0.01, ***^/###^p<0.001, ****^/####^p<0.0001. Hashsigns indicate comparison to WT, asterisks indicate comparison to mutant vehicle. Scale bar, 100 μm. n =6-8 animals/treatment.

### Short-term treatment with AF38469 improved histopathological outcomes in a CLN3 disease mouse model

We next sought to explore the effects of sortilin through AF38469 in another mouse model of an LSD with vastly different etiology, CLN3 Batten disease, which is characterized by juvenile onset and protracted progression. The *Cln3^Δex^*^7^*^/^*^8^ mouse model recapitulates the most common human pathogenic variant in CLN3 disease, leading to an absence of the transmembrane endosomal/lysosomal CLN3 protein, a putative lysosomal glycerophosphodiester transporter.^80, 81^ *Cln3^Δex^*^7^*^/8^*mice consistently exhibit progressive lysosomal pathology and neuroinflammation but have variable (i.e., inconsistent) lifespan and behavioral phenotypes.^68, 82–84^

Homozygous *Cln3^Δex^*^7^*^/^*^8^ and wild type mice received continuous treatment with AF38469 or vehicle via drinking water starting at wean until 16 weeks of age (**Figure 4A**), a time point when initial pathology is present but limited to no behavioral changes are apparent. As with the *Cln2^R^*^207^*^X^*study, we first tested the higher doses prevalent in the literature and saw a reduction in disease pathology with only the lower dose, encouraging us to move forward with a new study focusing on 10-fold and 100-fold lower doses (**Supplemental Figure 5A**). In this second lower dose study, we again observed profound improvement in a range of biochemical and histopathological measures.

AF38469-treated *Cln3^Δex^*^7^*^/^*^8^ mice exhibited an improvement of lysosomal enzyme activity as evidenced by increased TPP1 enzyme activity in brain lysates (**Figure 4B, C**). There was a significantly and potently decreased accumulation of lysosomal storage material (SubC) and microglial activation (CD68^+^) in the S1BF of the somatosensory cortex (**Figure 4D**), CA3 region of the hippocampus (**Figure 4E**), VPM/VPL of the thalamus (**Figure 4F**), and the dentate gyrus and hilus regions of the hippocampus (**Figure 4G**). Treatment with AF38469 significantly decreased astroglial activation (GFAP^+^) in the S1BF of the somatosensory cortex (low dose), CA3 region of the hippocampus (high dose), dentate gyrus and hilus regions of the hippocampus and the VPM/VPL of the thalamus (both doses) (**Figure 4D-G**). Treatment with AF38469 had no impact on the body weights of *Cln3^Δex^*^7^*^/^*^8^ mice, which are comparable to wild type (**Supplemental Figure 7B**). Given *Cln3^Δex^*^7^*^/^*^8^ mice have limited behavior changes at this early age, behavioral metrics were not measured.

### Short-term treatment with AF38469 reduces brain lysosomal storage and microglial activation in wild type mice

To examine whether sortilin inhibition could enhance lysosomal function and reduce neuroinflammation outside of the context of severe monogenic disease, we monitored the effects of AF38469 treatment on wild type mice. Wild type mice received continuous treatment with AF38469 or vehicle via drinking water starting at wean until 16 weeks of age. While higher doses (3 μg/ml, 78 μg/ml) of AF38469 had no impact of histopathological outcomes in wild type mice (**Supplemental Figure 6A**), treatment with the lower doses of AF38469 reduced lysosomal storage and microglial activation. The lower doses significantly decreased microglial activation (CD68^+^) in the S1BF of the somatosensory cortex and VPM/VPL of the thalamus. The lowest dose (3 μg/ml) of AF38469 significantly reduced the levels of astroglial activation (GFAP^+^) in the VPM/VPL of the thalamus of the AF38469 treated mice (**Supplemental Figure 6B**). Treatment with AF38469 (0.3 μg/ml) significantly increased the TPP1 enzyme activity in brain lysates (**Supplemental Figure 6D**). Treatment with AF38469 had no impact on the body weight of wild type mice (**Supplemental Figure 7D**). Collectively, these results demonstrate that sortilin inhibition enhances lysosomal function, reduces lysosomal storage material burden, and reduces markers of neuroinflammation even in the absence of severe disease.

## Discussion

Lysosomal dysfunction lies at the heart of a wide variety of disease conditions, ranging from the rare and often severe monogenic LSDs, to more common disorders such as Parkinson’s disease, frontotemporal dementia, and aging associated decline.^85^ Therapies that can boost lysosomal function thus have vast potential for enhancing human health. Preservation or restoration of lysosomal dysfunction in disease contexts can restore cellular recycling and energy maintenance pathways, disrupt downstream pathogenic cascades, and prevent the functional deterioration or death of affected cells. Ideally, such a therapy would enhance lysosomal function in diverse cell types in a disease agnostic manner, even after the onset of cellular pathology.^86^ Here, we demonstrate that sortilin inhibition fits these criteria, potentially offering utility for the treatment of a wide variety of conditions characterized by lysosomal dysfunction.

Given the roles of sortilin in lysosomal biogenesis, complete inhibition of this pathway might be expected to produce a deleterious response. On the other hand, incomplete inhibition could activate a beneficial compensatory response, as has been shown for many other drugs that exhibit the phenomenon of hormesis.^58^Indeed, we found that low-level inhibition of SORT1 enhanced lysosomal gene transcription via activation and nuclear translocation of TFE3 and TFEB, master regulators of lysomosal biogenesis through the CLEAR network.^36^ This response enhanced lysosomal enzyme activity as reflected by increased activities for PPT1 and TPP1 and reduced storage material burden as evidenced by reductions in ASM and SubC. In animal models of two LSDs, this rescue of lysosomal health ameliorated neuroinflammation as evidenced by reductions in CD68 and GFAP immunoreactivity.

The precise molecular pathways by which this activity is manifested have not yet been elucidated. SORT1 and the SORT1 ligand PGRN have both been shown to activate mTORC1, a critical inhibitor of TFEB and TFE3 translocation and function.^87, 88^ Low-level SORT1 inhibition could reduce PGRN binding and subsequently mTORC1 activity, disinhibiting TFEB and TFE3 without interrupting the lysosomal enzyme sorting functions of SORT1. Such a mechanism would be consistent with a dependence on the normal functioning of the PGRN/SORT1 axis, though as our results showed an increase in enzyme activity in *PGRN^-/-^* cells, it is clear that AF38469 does not necessarily work entirely through PGRN. Therefore future studies should examine the precise mechanisms by which low-level SORT1 inhibition has benefits for lysosomal function outside of this known mechanism.

Regardless of the precise mechanisms, our studies support SORT1 as a promising new drug target for lysosomal storage disorders. In two mouse models of neurocentric LSDs, lysosomal function and health was improved upon treatment with AF38469. Treatment with low doses of AF38469 prevented the acculumation of histopathological markers associated with CLN2 and CLN3 disease. Most strikingly, microglial activation was reduced to near wild type levels in the thalamus, a brain region that has been refractory to rescue with existing gene therapy drug candidates.

This work was focused on neurodegenerative LSDs; however, given the ubiquitous expression of sortilin throughout peripheral tissues, SORT1 inhibition could have value as a therapeutic for other LSDs with primary complications in other body systems such as Fabry, Pompe, or Gaucher disease. Furthermore, SORT1 may be a druggable target for other diseases with a lysosomal component, such as Parkinson’s disease or even aging. Since the efficacious doses for lysosomal benefits are orders of magnitude lower than those used to modulate other SORT1-dependent functions,^69, 90–92^ it is likely that beneficial responses could be activated without interrupting normal homeostatic functions.

## Authors Contributions

Conceptualization: TBJ, JJB, JMW; Methodology: CD, MAP, CDB, TBJ, JJB, JMW; JMW; Formal Analysis: HGL, JTA, KJT, CD, MAP, KAW, MJR, BLM, JJB; Investigation: HGL, JTA, KJT, CD, MAP, CDB, MJR, BLM; Writing – Original Draft: HGL, JJB; Writing: Review & Editing: HGL, JTA, KJT, CD, MAP, CDB, KAW, MJR, BLM, TBJ, JJB, JMW; Visualization: HGL, JTA, KJT, CD, MAP; Supervision and Project Administration: MAP, CDB, KAW, TBJ, JJB, JMW; Funding Acquisition: JMW.

## Acknowledgements

This work was supported by funding to JMW from the Forebatten Foundation and Lehrman Family Fund. This work also received support from the Sanford Research Histology and Imaging Core within the Sanford Research Center for Pediatric Research and Cancer Biology (NIH P20GM103620 and P20GM103548) and the Sanford Research Design and Biostatistics Core within the Sanford Research Center for Health Outcomes and Population Research (NIH P20GM121341). GRN (*Cln11-/-*) knockout fibroblasts were a gift from the Kukar lab at Emory University.

## Materials and Methods

### Ethics Statement/Animals

All animal studies were housed under identical conditions in an AAALAC International-accredited facility with Institutional Animal Care and Use Committee (IACUC) approval (US Department of Agriculture [USDA] license 46-R-0009; protocol #178-02-24D, Sanford Research, Sioux Falls, SD, USA). Wild type, *Cln1^R^*^151^*^X^* (JAX #026197, *Cln2^R^*^207^*^X^* (JAX #030696), *Cln3^Δex^*^7^*^/^*^8^ (JAX #017895), *Cln6^nclf^* (JAX #003605), and *Cln8^mnd^* (JAX #003906) mutant mice on C57BL/6J backgrounds were used for all studies For behavior studies and immunohistochemistry experiments, an n of 7-8 and 4-8 mice, respectively, per group were used (**Table 1 and 2**).

#### Primary neuronal cells (PNC) generation, treatment, and imaging

On embryonic day (E) 15.5, mouse cortical neurons were prepared as previously described^93^. Briefly, the cortices were collected and dissociated in 0.32 mg/ml L-Cysteine hydrochloride, 19.5 units/ml Papain, and 37.5 µg/ml DNase. The cells were resuspended in neurobasal medium supplemented with B27 (1X) and L-glutamine (2 mM). The cells were cultured at 5×10^5^ cells/ml (100 µL/well, 96-well plate) coated with a Laminin (8.33 µg/ml) and Poly-D-Lysine (83.3 µg/ml) solution and placed in an incubator (37°C, 5% CO_2_).On *in vitro* day (DIV) 3 and DIV5, cells were treated with AF38469 (40 nM, 400 nM, 4 μM) or DMSO vehicle along with a 50% medium change using supplemented Neurobasal medium. On DIV7, the well medium was replaced with the following dyes dissolved in unsupplemented Neurobasal medium: Hoechst 33342 (Thermo Fisher, H3570, 1:1000 dilution) and LysoTracker™ Red DND-99 (Thermo Fisher, L7528, 1:10,000 dilution). Following a 20-minute incubation (37°C, 5% CO_2_), the cells were incubated with a fixative solution (2% sucrose, 2% paraformaldehyde (PFA)) in PBS for five minutes. The plate was imaged in PBS and analyzed using CellInsight CX7 High-Content Screening Platform (CX7) and associated software.

#### Mouse embryonic fibroblasts (MEF) generation, treatment, and imaging

On E15.5, MEFs were isolated as previously described with minor modifications^94^. Briefly, following the inhibition of trypsin with Dulbecco’s Modified Eagle Medium (DMEM) with High Glucose, 4.0 mM L-Glutamine, and Sodium Pyruvate supplemented with 10% Fetal Bovine Serum (FBS) and 1% Penicillin-Streptomycin (MEF medium), the cell mixture was filtered through a 70 µm filter and centrifuged. The cells were resuspended in MEF medium, plated in 10 cm dishes, and placed in an incubator (37°C, 5% CO_2_). For assays, cells were resuspended as above and plated at 3.5e4 cells/ml (100 µl/well, 96-well plate). On DIV3 and DIV5, cells were treated with AF38469 (40 nM, 400 nM, 4 μM) or DMSO control along with a 100% medium change using MEF medium (3% FBS). On DIV7, the well medium was replaced with the dyes descibed above dissolved in unsupplemented MEF medium. Following a 20-minute incubation (37°C, 5% CO_2_), the cells were incubated with a fixative solution (2% sucrose, 2% PFA) in PBS for five minutes. The plate was imaged in PBS and analyzed using CellInsight CX7 High-Content Screening Platform (CX7) and associated software.

#### *In vitro* comparative transcriptomics

Wild type MEFS were prepared as described above.The cells were plated at 7×10^4^ cells/ml with a volume of 10 ml/dish in 10 cm dishes. On DIV3, the cells were treated with AF34869 (40 nM) along with a 100% medium change using DMEM with High Glucose, 4.0 mM L-Glutamine, and Sodium Pyruvate supplemented with 3% Fetal Bovine Serum and 1% Penicillin-Streptomycin. On DIV5, total RNA was extracted using Qiagen RNeasy® Mini Kit (Qiagen, 74004) according to the manufacturer’s instructions. RNA sequencing was performed by Novogene. Gene ontology analysis and pathway analysis was performed using the DAVID functional annotation tool (https://david.ncifcrf.gov/summary.jsp). The official gene symbol and Mus musculus were selected as the identifier and the species of origin in this tool. Threshold for the number of genes per term was set to 2 and EASE threshold set to 0.1

#### *In vitro l*ysosomal enzyme activity assays

MEFs were prepared and treated as described above. The cells were plated at 5.3×10^4^ cells/ml (2 ml/well, 6-well plate). On DIV7, the cells were manually disassociated. Enzyme activities of TPP1 (tripeptidyl peptidase 1) and PPT1 (palmitoyl protein transferase 1) were evaluated as previously described with minor modifications^95^. Protein quantification using the Pierce 660 nm Protein Assay (Thermo Fisher, 22660) was performed. Assays were performed in 96 well plates with 10 µg of protein sample per well. The fluorescence was evaluated using BioTek Cytation 3 (PPT1: Excite 355 nm/Emit 460 nm, TPP1: Excite 380 nm/Emit 460 nm).

#### Transfection for TFEB/TFE3 and Live Imaging

Neuro2a cells were plated at 1.5×10^5^ cells/ml (1 ml per well, 24-well plate) using DMEM with High Glucose, 4.0 mM L-Glutamine, and Sodium Pyruvate supplemented with 3% Fetal Bovine Serum and 1% Penicillin-Streptomycin. pEGFP-N1-TFE3 (Addgene, 3812) and pEGFP-N1-TFEB (Addgene, 38119) were transfected into Neuro2a cells using Lipofectamine 3000 (Thermo Fisher, L3000008) transfection following the manufacturer’s provided protocol (Fisher #L3000015). After a 24-hour incubation, the cells received a 100% medium change.

### For nuclear translocation analysis with live imaging

On DIV3, the cells were incubated with Hoechst (Thermo Fisher, H3570, 1:1000) for 20 minutes (37°C, 5% CO_2_) and the plate was imaged and analyzed using CellInsight CX7 High-Content Screening Platform (CX7) and associated software. Following imaging, cells were treated with AF38469 (40 nM and 400 nM) or DMSO with 100% medium change. The compounds were incubated for 3 hours (37°C, 5% CO_2_), with imaging and analysis using CellInsight CX7 High-Content Screening Platform (CX7) and associated software every 30 minutes during that timeframe. Between imaging sessions, the plate was kept in an incubator (37°C, 5% CO_2_).

### For nuclear translocation analysis with immunocytochemistry

On DIV3, a subset of cells was incubated for 20 minutes at room temperature with fixative solution containing 2% sucrose and 2% PFA in PBS. A second subset of cells was treated with AF38469 (40 nM and 400 nM) or DMSO with a 100% medium change. The compounds were incubated for 3 hours (37°C, 5% CO_2_). After compound incubation, fixative solution was added and the plate was processed for immunocytochemistry as described below. Cells were incubated with blocking buffer (PBS, 0.1% Saponin, 15% Serum) for 20 minutes at room temperature with light agitation. The cells were incubated overnight at 4°C on agitator with anti-GFP (Abcam, ab13970; 1:1000) in tris-buffered saline 1X (TBS) with 10% fetal bovine serum. Cells were incubated for 2 hours at room temperature with goat anti-chicken IgY (H+L) Alexa Fluor™ 488 (Invitrogen, A-11039, 1:1000) diluted in TBS with 10% Serum. Coverslips were mounted on glass slides using aqueous mounting medium and imaged using the Nikon A1R Ti2E Confocal and associated Nikon software. For immunocytochemistry analysis, the slides were imaged on Nikon NiE microscope. The files were saved in the Nikon Elements .ND2 format and were converted into 8-bit mono .tifs for each channel. The images were analyzed using ImageJ (version 1.52). In brief, each individual image was subdivided into four non overlapping fields, and image threshold analysis and image particle analysis macros were used.

#### Tissue collection and preparation

Wild type, *Cln3*^Δex7/8^, and *Cln2^R^*^207^*^X^* mutant mice were CO_2_ euthanized at 16 and 11 weeks of age, respectively, and perfused with PBS. For enzyme and transcriptomic analysis, brain tissue was flash frozen with 2-methylbutane and stored at -80°C. Tissues for enzyme activity were homogenized in protein lysis buffer [50 mM Tris-HCl pH 7.4; 150 mM NaCl; 0.2% Triton X-100; 0.3% NP-40; 0.1 mM PMSF; 1x HALT Protease Inhibitor Cocktail (Thermo Fisher Scientific, 78429)] and TPP1 and PPT1 enzyme activities were completed as described above. For immunohistology analysis,brains were fixed in 4% PFA for 24 hours then stored at 4°C in 0.02% sodium azide until futher use.

#### Immunohistochemistry

Immunohistochemistry was performed as previously described with minor modifications^95^. Briefly, fixed brains were embedded in 3% agarose and sectioned at 50 μm thickness with a vibrating microtome (Leica VT1000S Microtome). Free floating sections were incubated in 1% hydrogen peroxide for 15 minutes then incubated with anti-ATP synthase subunit C (Abcam, ab181243; 1:1000), anti-GFAP (Dako, Z0334; 1:8000), or anti-CD68 (AbD Serotec, MCA1957; 1:2000),diluted in 15% goat serum-containing TBS-T (TBS with 0.3% Triton X-100) overnight at 4°C. Tissues were incubated with appropriate secondary antibodies: goat anti-rat (Vector Labs, BA-9400 1:1000) or goat anti-rabbit biotinylated (Vector Labs, BA-1000 1:1000) diluted in TBS-T+ 10% goat serum and followed by incubation with VECTASTAIN^®^ Elite^®^ ABC-HRP Kit, Peroxidase (Standard) (Vector Labs, PK-6100). Immunoreactivity was visualized using a 0.05% 3,3—Diaminobenzidine (DAB) reaction (Sigma Aldrich, D5905-50TAB). For immunohistochemistry analysis, slides were scanned on a Aperio AT2 Slide Scanner (AN7621) at 20X and were visualized Aperio ImageScope x64 (12.04.02.5010). Labeling intensities for both colorimetric immunohistochemical preparations were quantified using ImageJ software (version 1.52)

#### Behavioral Testing

*Cln2^R^*^207*X*^ mice treated with 250, 10, 1, and 0.1 μg AF38469 (n= 8,8,8, 7 respectively), *Cln2^R^*^207*X*^ mice treated with DMSO vehicle control (n=7), wild type mice treated with 250, 10, 1, and 0.1 μg AF38469 (n= 8,8,7, 7 respectively), and wild type mice treated with DMSO control (n= 8) were tested at 11 weeks of age as previously described over the course of 20 minutes using a force plate actimeter (BASi, West Lafayette, IN)^53^. Tremor scores (5-10 Hz, 10-15 Hz, 15-20 Hz, 20-25 Hz) were analyzed.

#### Statistical Analysis

Statistical analyses were performed with GraphPad Prism (v8 or later) as previously described^96^. Specific details are listed in the figure legends. Detailed sample N’s are shown in **Table 1 and 2**. For *in vitro* experiments, the number of cells analzyed/well was a range between 1500-2000, and three wells/condition were analyzed as technical replicated; a one-way ANOVA with a Šidák post-hoc test was performed comparing each group with the wild type untreated and the corresponding mutant vehicle treated. For *in vivo* histopathological quantitation, 3 brain slices from each animal were used for each assay. Four images /region/tissue were taken equating to 12 technical replicates/region/animal. A nested one-way ANOVA with a Šidák post-hoc test was performed comparing each group with the wild type untreated and the corresponding mutant vehicle treated. For behavior testing, 7-8 animals were tested, and a one-way ANOVA with a Šidák post-hoc test was preformed comparing each group with the wild type untreated and the corresponding mutant vehicle treated .Statistical outliers, as determined by GraphPad Prism, and obvious techincal failures (e.g., loss of desired region during processing, tissue folding, bubbles) were removed which accounts for any variability in technical replicates in **Table 1 and 2**.

