## Supplemental Figures for "Sortilin inhibition treats multiple neurodegenerative lysosomal storage disorders"

**Supplemental Figure 1 - Inhibition of sortilin through ART1001 (40 nM, 400 nM, 4µM) impacts the accumulation of ASM and Lysotracker in cell models of Batten disease.** Mouse embryonic fibroblasts (MEFs) were isolated from *Cln1*$^{R151X}$, *Cln2*$^{R207X}$, *Cln3*$^{\Delta ex7/8}$, *Cln6*$^{nclf}$, and *Cln8*$^{mnd}$ mouse lines. *Cln11*$^{-/-}$ fibroblasts were isolated from adult *Cln11*$^{-/-}$ mice All cell lines were dosed with drug-containing media on DIV3 and DIV5 and analyzed on DIV7 using the CellInsight CX7 High-Content Screening Platform (CX7) (A-F) Upon treatment with ART1001 (40 nM, 400 nM, 4 µM) ASM accumulation was reduced in WT, *Cln1*$^{R151X}$ *Cln2*$^{R207X}$, *Cln3*$^{\Delta ex7/8}$, *Cln6*$^{nclf}$, and *Cln8*$^{mnd}$ MEFs and (G) increased in *Cln11*$^{-/-}$ MEFs. (H-J, L-M) Treatment with ART1001 (40 nM, 400 nM, 4 µM) decreased the Lysotracker accumulation in WT, *Cln1*$^{R151X}$, *Cln2*$^{R207X}$, *Cln6*$^{nclf}$, *Cln8*$^{mnd}$, and *Cln11*$^{-/-}$ MEFs. (K) Lysotracker signal was increased in *Cln3*$^{\Delta ex7/8}$ MEFs upon treatment with ART1001 (40 nM) and decreased when cells were treated with 400 nM and 4 µM doses (O-U) Treatment with ART1001 (40 nM, 400 nM, 4 µM) had no impact on cellular viability compared to vehicle- treated mutant (S) aside from *Cln6*$^{nclf}$ (400 nM) which was reduced. (A-M). Mean ± S.E.M. of % of the total cell area. (O-U) Mean ± S.E.M. of number of cells reflected as valid object count. One-way ANOVA with a Šidák post-hoc test $^{*/\#}p<0.05$, $^{**/\#\#}p<0.01$, $^{***/\#\#\#}p<0.001$, $^{****/\#\#\#\#}p<0.0001$. Hashsigns indicate comparison to WT, asterisks indicate comparison to mutant vehicle. (A-M) n = 650 – 4500 cells/treatment (O-U) n = 9 wells/treatment. Scale bar, 100 µm.

**Supplemental Figure 2 - Inhibition of sortilin through ART1001 (40 nM, 400 nM, 4µM) impacted the accumulation of ASM and Lysotracker in neuronal cell models of Batten disease**. Primary neuronal cells (PNCs) isolated from *Cln1*$^{R151X}$, *Cln2*$^{R207X}$, *Cln3*$^{\Delta ex7/8}$, *Cln6*$^{nclf}$, and *Cln8*$^{mnd}$ mouse lines. All cell lines were dosed with drug-containing media on DIV3 and DIV5 and analyzed on DIV7 using the CellInsight CX7 High-Content Screening Platform (CX7) (A-F) Upon treatment with ART1001 (40 nM, 400 nM, 4µM), ASM accumulation was generally reduced in WT, *Cln1*$^{R151X}$ *Cln2*$^{R207X}$, *Cln3*$^{\Delta ex7/8}$, and *Cln8*$^{mnd}$ PNCs; and (E) increased in *Cln6*$^{nclf}$ PNCs (b)

Upon treatment with ART1001 (40 nM, 400 nM, 4μM), Lysotracker accumulation was reduced in WT, *Cln1*$^{R151X}$, *Cln6*$^{nclf}$ PNCs, and was variably impacted in *Cln2*$^{R207X}$, *Cln3*$^{\Delta ex7/8}$, and *Cln8*$^{mnd}$ PNCs. (G-L) Treatment with ART1001 (40 nM, 400 nM, 4μM) had no impact on cellular viability compared to vehicle-treated mutant. (A-L). Mean ± S.E.M. of % of the total cell area. (M-R) Mean ± S.E.M. of number of cells reflected as valid object count. One-way ANOVA with a Šidák post-hoc test $^{*/\#}p<0.05$, $^{**/\#\#}p<0.01$, $^{***/\#\#\#}p<0.001$, $^{****/\#\#\#\#}p<0.0001$. Hashsigns indicate comparison to WT, asterisks indicate comparison to mutant vehicle. (A-L) n = 4000 - 20000 cells/ treatment (M-R) n = 9 wells/treatment. Scale bar, 100 μm, Inset scale bar, 50 μm.

**Figure 3 - Sortilin inhibition through ART1001 treatment increases enzyme activity of PPT1 and TPP1 *in cellulo.*** Mouse embryonic fibroblasts (MEFs) were isolated from WT, *Cln2*$^{R207X}$, *Cln3*$^{\Delta ex7/8}$, and *Cln6*$^{nclf}$ mouse embryos, and *Cln11*$^{-/-}$ fibroblasts were isolated from adult *Cln11*$^{-/-}$ mice. Cells were cultured and treated with ART1001 (40nM) on DIV3 and DIV5; the cells were collected on DIV7, and the enzyme activity assays were performed on the cellular lysates. (A-E) Treatment with ART1001 increased PPT1 enzyme activity in all cell lines. (G-J) Treatment with ART1001 increased TPP1 enzyme activity in all cell lines except for (F) *Cln2*$^{R207X}$ MEFs. Mean± S.E.M. One-way ANOVA with a Šidák post-hoc test *p <0.05, **p<0.01, ***p<0.001, ****p<0.0001. n = 3-5 wells/treatment.

**Supplemental Figure 4 - Inhibition of sortilin through short-term treatment with ART1001 impacted histopathological and behavioral outcomes in *Cln2*$^{R207X}$ mice.** Homozygous *Cln2*$^{R207X}$ and litter mate wild type mice received continuous treatment with ART1001 or vehicle via the drinking water starting at wean until 11 weeks of age. Body weights were measured biweekly until 10 weeks of age when force plate measurements were collected. (A) ART1001 treatment (78 µg/ml) in *Cln2*$^{R207X}$ mice prevented SubC accumulation and had no impact on microglial reactivity (CD68$^{+}$) or astroglial activation (GFAP$^{+}$) in the S1BF of the somatosensory cortex (B) ART1001 treatment (78 µg/ml) in *Cln2*$^{R207X}$ mice had no impact of accumulation of mitochondrial ATP synthase subunit C ($SubC^+$), microgliosis ($CD68^+$), or astrocytosis ($GFAP^+$) in the VPM/VPL of the thalamus. Mean ± S.E.M. Nested one-way ANOVA with a Šidák post-hoc test $^{*/\#}p<0.05$, $^{**/\#\#}p<0.01$, $^{***/\#\#\#}p<0.001$, $^{****/\#\#\#\#}p<0.0001$. Hashsigns indicate comparison to WT, asterisks indicate comparison to mutant vehicle. n = 7-8 animals/treatment. Scale bar, 100 μm.

**Supplemental Figure 5 - Inhibition of sortilin through short-term treatment with ART1001 impacted histopathological in *Cln3*$^{\Delta ex7/8}$ mice.** Homozygous *Cln3*$^{\Delta ex7/8}$ and wild type mice received continuous treatment with ART1001 or vehicle via drinking water starting at wean until 16 weeks of age. Body weights were measured monthly. (A ART1001 treatment (3 μg/ml, 78 μg/ml) in *Cln3*$^{\Delta ex7/8}$ mice prevented SubC accumulation and had no impact on microglial reactivity ($CD68^+$) or astroglial activation ($GFAP^+$) in the S1BF of the somatosensory cortex. (B) ART1001 treatment (3 μg/ml) prevented accumulation of mitochondrial ATP synthase subunit C ($SubC^+$) in the VPM/VPL nuclei of the thalamus (3 μg/ml) when compared to vehicle treated mice. Mean ± S.E.M. Nested one-way ANOVA with a Šidák post-hoc test $^{*/\#}p<0.05$, $^{**/\#\#}p<0.01$, $^{***/\#\#\#}p<0.001$, $^{****/\#\#\#\#}p<0.0001$. Hashsigns indicate comparison to WT, asterisks indicate comparison to mutant vehicle. n = 6-8 animals/treatment. Scale bar, 100 μm.

**Supplemental Figure 6 - Inhibition of sortilin through short-term treatment with ART1001 impacted histopathological and behavioral outcomes in wild type mice**. (A) Wild type mice received continuous treatment with ART1001 or vehicle via drinking water starting at wean until 16 weeks of age. Body weights were measured monthly. (B) Treatment with ART1001 had no impact on PPT1 enzyme activity levels in WT treated mice (C) Treatment with ART1001 increased TPP1 enzyme activity levels in WT treated mice (D-E) ART1001 treatment (3 μg/ml, 78 μg/ml) in WT mice had no impact of accumulation of mitochondrial ATP synthase subunit C ($SubC^+$), microgliosis ($CD68^+$), or astrocytosis ($GFAP^+$) in the S1BF of the somatosensory cortex or the VPM/VPL of the thalamus. (F) Treatment with ART1001 reduced microgliosis ($CD68^+$) and had no impact on SubC accumulation or astrocytosis ($GFAP^{+}$) in the S1BF of the somatosensory cortex. (G) Treatment with ART1001 reduced SubC accumulation, microgliosis ($CD68^{+}$), and astrocytosis ($GFAP^{+}$) in the VPM/CVPL of the thalamus. Mean ± S.E.M. Nested one-way ANOVA with a Šidák post-hoc test $^{*/\#}p<0.05$, $^{**/\#\#}p<0.01$, $^{***/\#\#\#}p<0.001$, $^{****/\#\#\#\#}p<0.0001$. Hashsigns indicate comparison to WT, asterisks indicate comparison to mutant vehicle. n = 6-8 animals/treatment. Scale bar, 100 μm.

**Supplemental Figure 8** Body weights measured at timepoints indicated along X axis. (A) ART1001 treated $Cln2^{R207X}$ mice (B) ART1001 treated $Cln3^{\Delta ex7/8}$ mice and (C) ART1001 treated wild type mice. Mean ± S.E.M. Two-way ANOVA with Dunnett's multiple comparisons test. *p <0.05, **p<0.01, ***p<0.001, ****p<0.0001 compared to WT. n = 1-4 sex/treatment

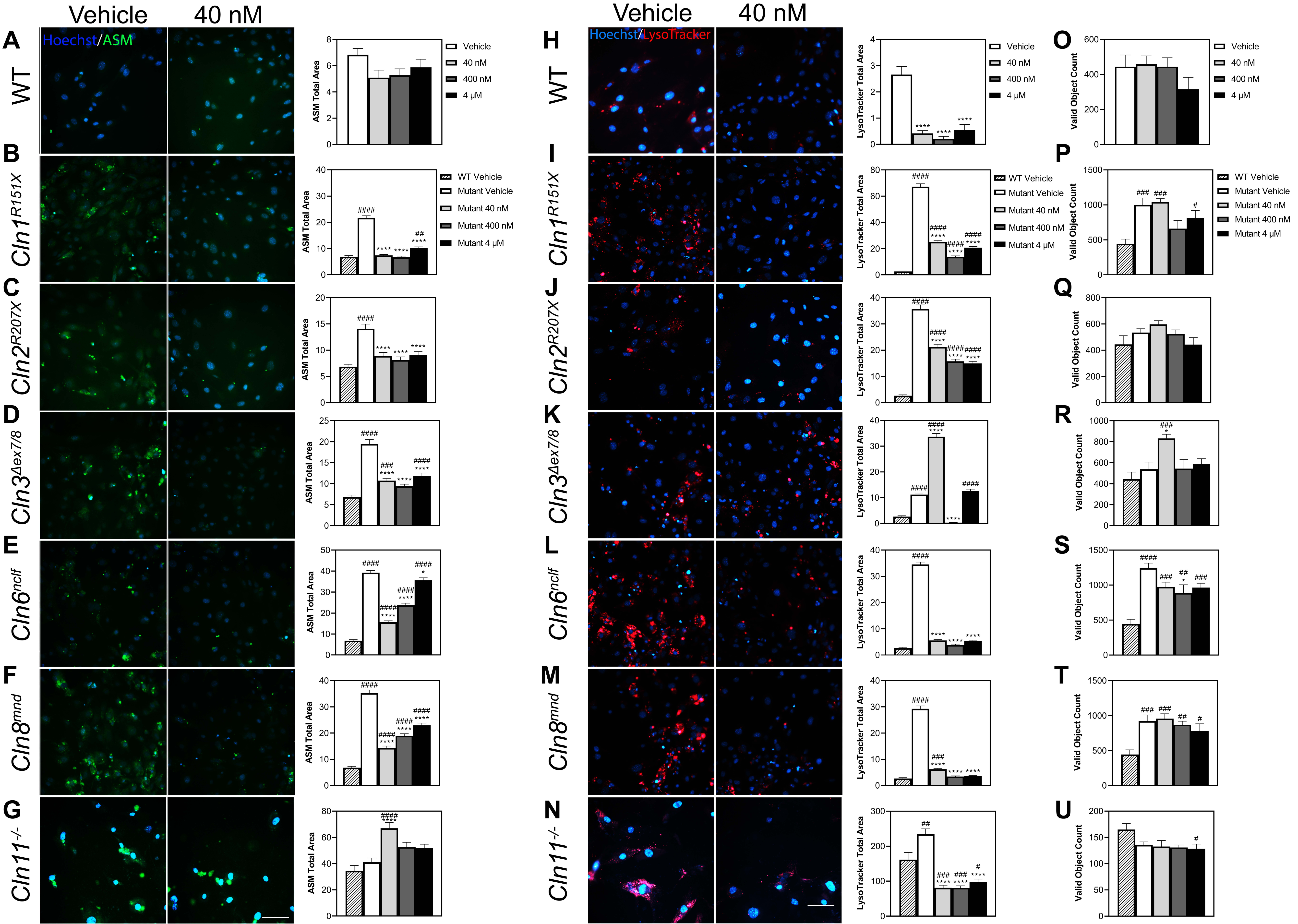

Vehicle
40 nM
Hoechst/ASM
Hoechst/LysoTracker
A
B
C
D
E
F
G
H
I
J
K
L
M
N
O
P
Q
R
S
T
U
WT
Cln1R151X
Cln2R207X
Cln3Δex7/8
Cln6nclf
Cln8mnd
Cln11-/-
ASM Total Area
LysoTracker Total Area
Valid Object Count
Vehicle
40 nM
400 nM
4 μM
WT Vehicle
Mutant Vehicle
Mutant 40 nM
Mutant 400 nM
Mutant 4 μM

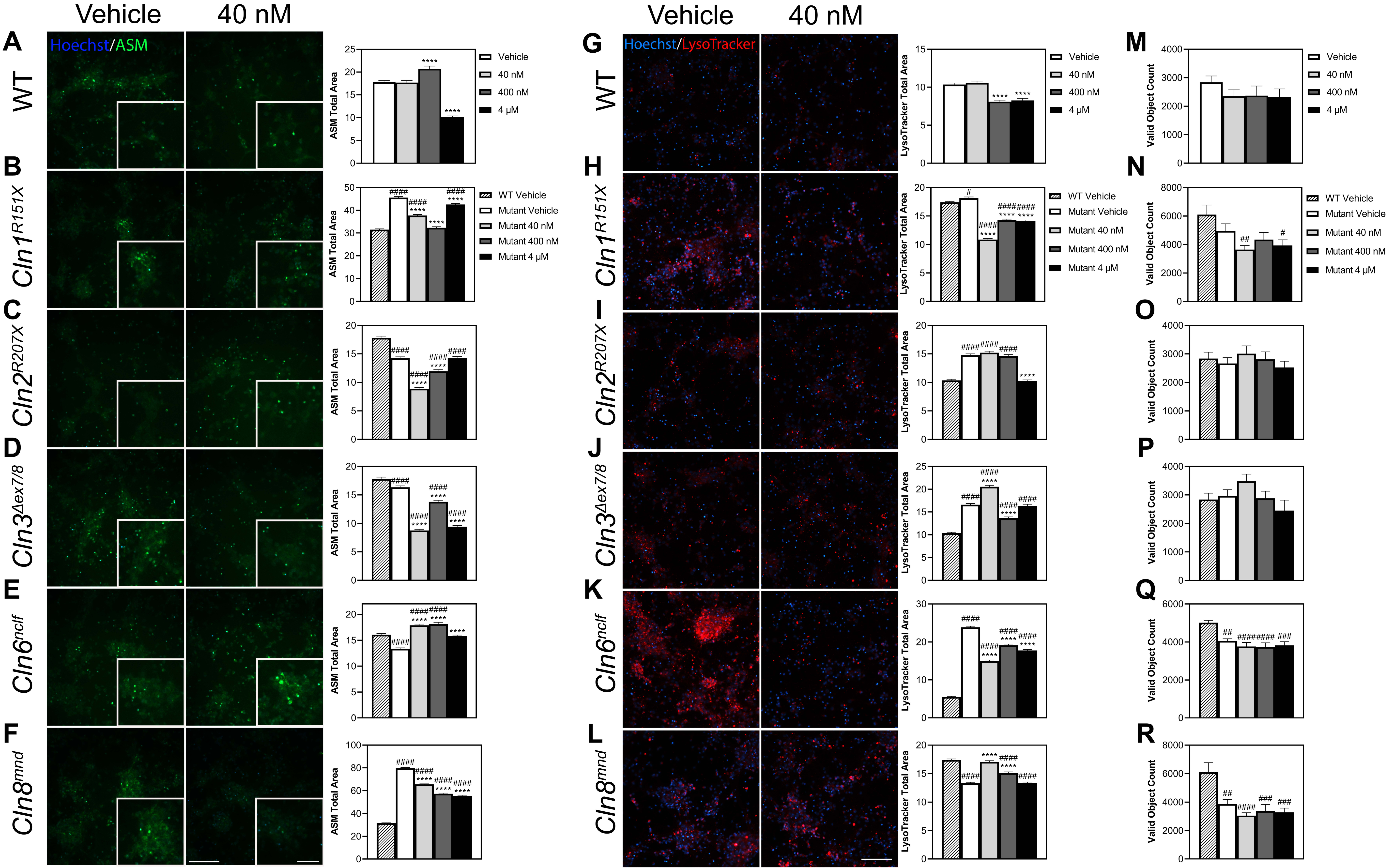

Vehicle
40 nM
Hoechst/ASM
Hoechst/LysoTracker
WT
Cln1R151X
Cln2R207X
Cln3Δex7/8
Cln6nclf
Cln8mnd
ASM Total Area
LysoTracker Total Area
Valid Object Count
Vehicle
40 nM
400 nM
4 μM
WT Vehicle
Mutant Vehicle
Mutant 40 nM
Mutant 400 nM
Mutant 4 μM

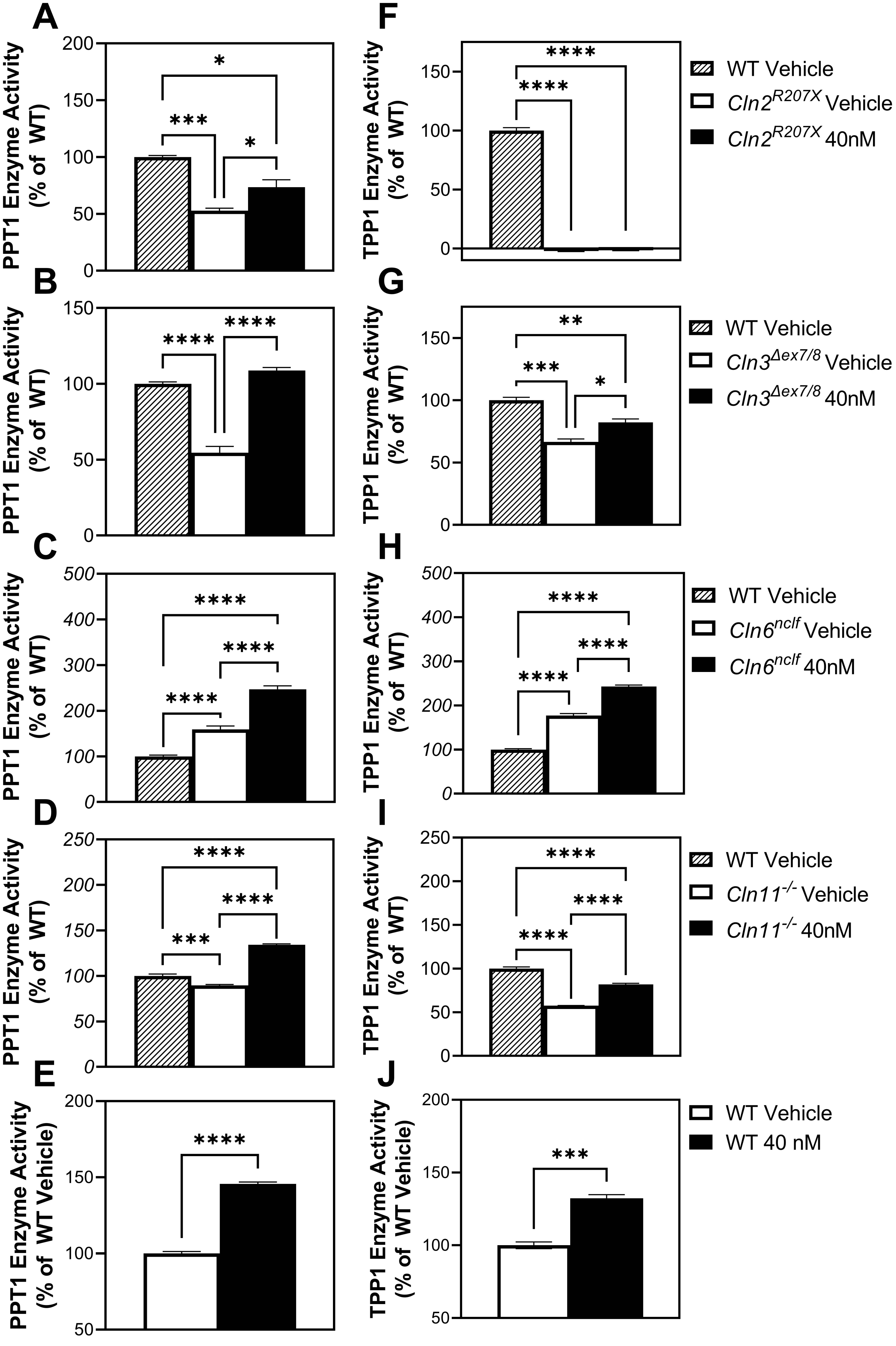

A
PPT1 Enzyme Activity (% of WT)
B
PPT1 Enzyme Activity (% of WT)
C
PPT1 Enzyme Activity (% of WT)
D
PPT1 Enzyme Activity (% of WT)
E
PPT1 Enzyme Activity (% of WT Vehicle)
F
TPP1 Enzyme Activity (% of WT)
WT Vehicle
Cln2R207X Vehicle
Cln2R207X 40nM
G
TPP1 Enzyme Activity (% of WT)
WT Vehicle
Cln3Δex7/8 Vehicle
Cln3Δex7/8 40nM
H
TPP1 Enzyme Activity (% of WT)
WT Vehicle
Cln6nclf Vehicle
Cln6nclf 40nM
I
TPP1 Enzyme Activity (% of WT)
WT Vehicle
Cln11-/- Vehicle
Cln11-/- 40nM
J
TPP1 Enzyme Activity (% of WT Vehicle)
WT Vehicle
WT 40 nM

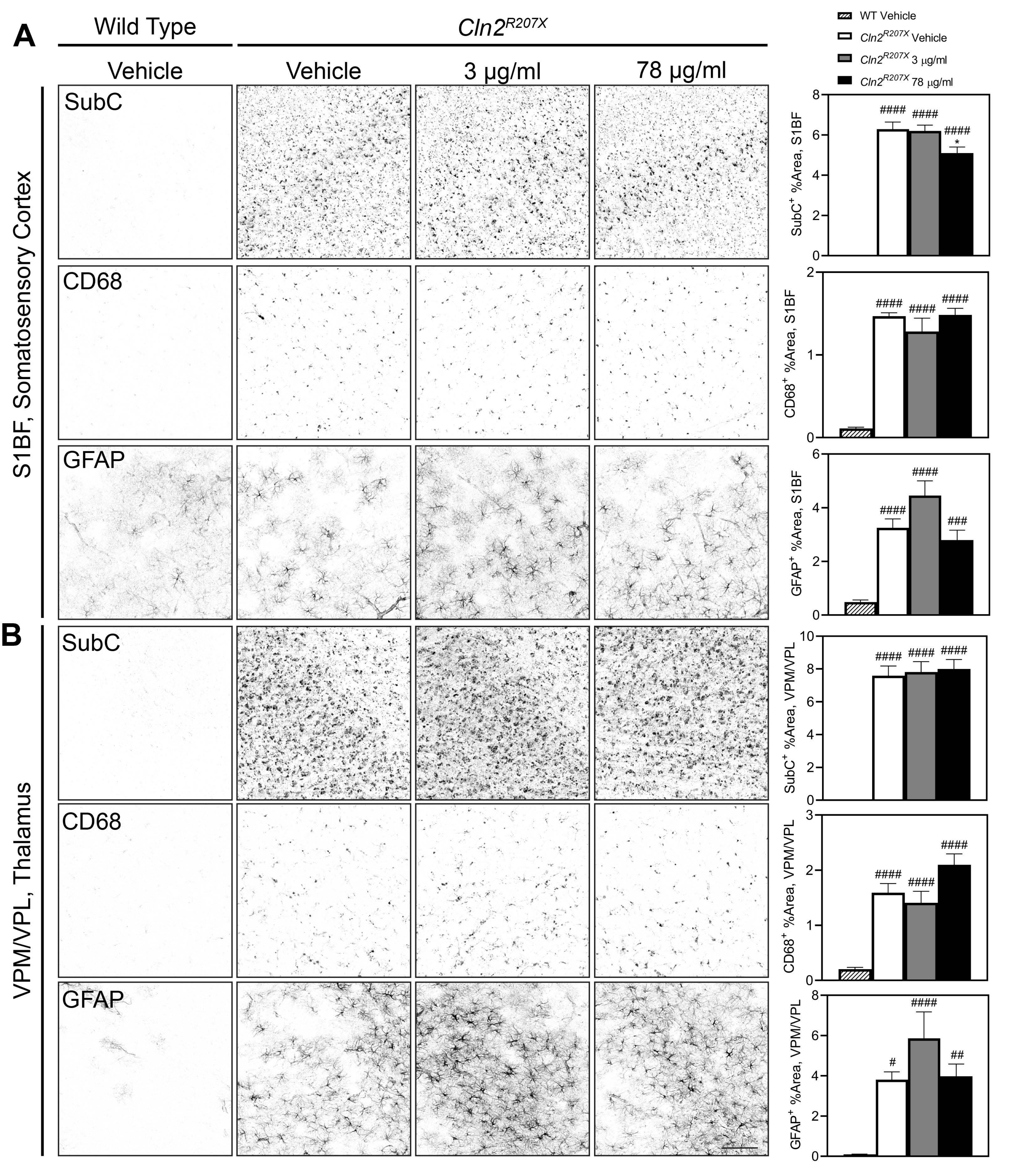

A
Wild Type
Cln2^R207X
Vehicle
Vehicle
3 µg/ml
78 µg/ml
S1BF, Somatosensory Cortex
SubC
CD68
GFAP
WT Vehicle
Cln2^R207X Vehicle
Cln2^R207X 3 µg/ml
Cln2^R207X 78 µg/ml
SubC+ %Area, S1BF
CD68+ %Area, S1BF
GFAP+ %Area, S1BF
####
###
*
B
VPM/VPL, Thalamus
SubC
CD68
GFAP
SubC+ %Area, VPM/VPL
CD68+ %Area, VPM/VPL
GFAP+ %Area, VPM/VPL
####
##
#

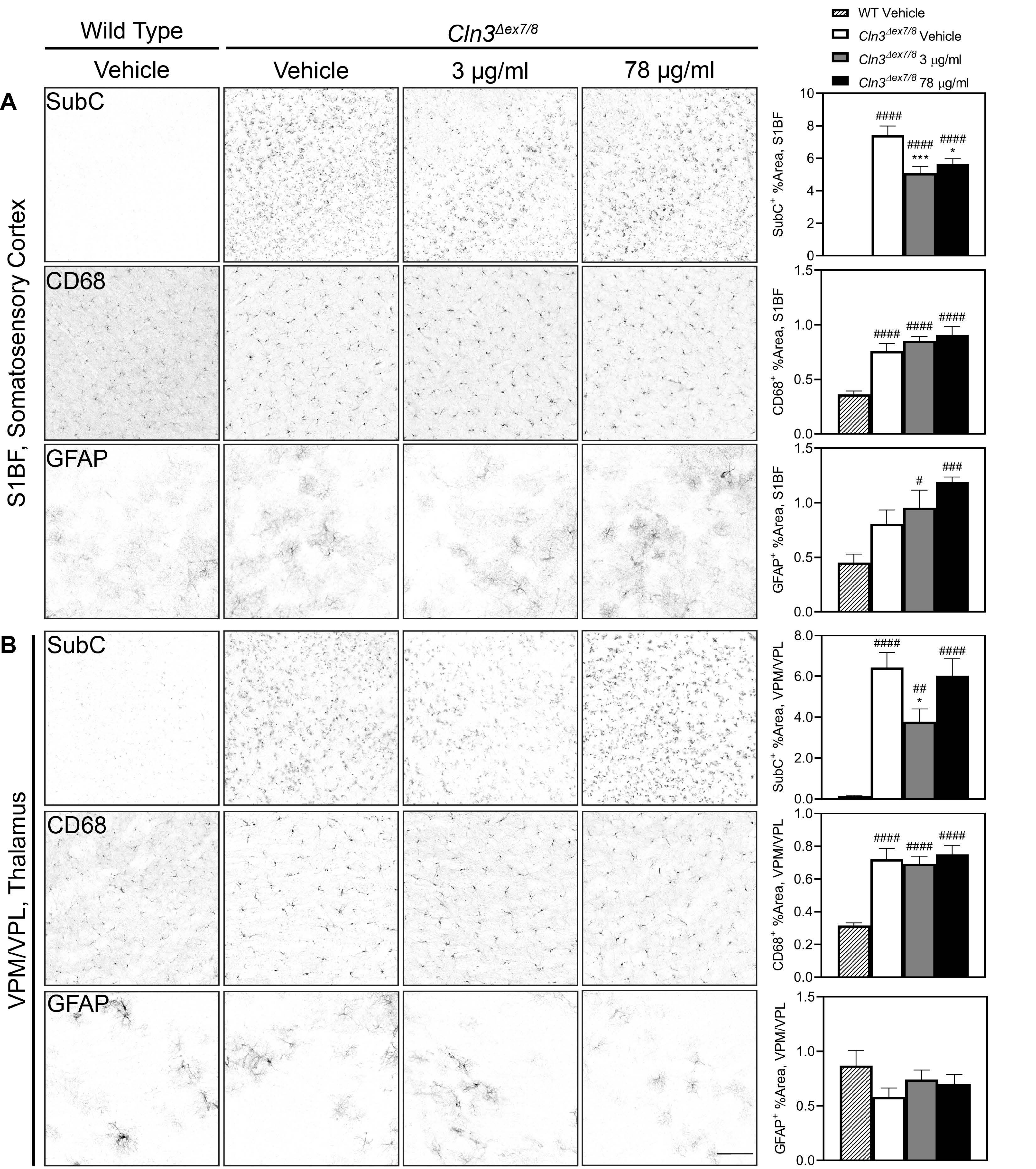

Wild Type
Cln3Δex7/8
Vehicle
Vehicle
3 μg/ml
78 μg/ml
A
S1BF, Somatosensory Cortex
SubC
CD68
GFAP
B
VPM/VPL, Thalamus
SubC
CD68
GFAP
WT Vehicle
Cln3Δex7/8 Vehicle
Cln3Δex7/8 3 μg/ml
Cln3Δex7/8 78 μg/ml
SubC+ %Area, S1BF
CD68+ %Area, S1BF
GFAP+ %Area, S1BF
SubC+ %Area, VPM/VPL
CD68+ %Area, VPM/VPL
GFAP+ %Area, VPM/VPL
####
***
*
##
#
###

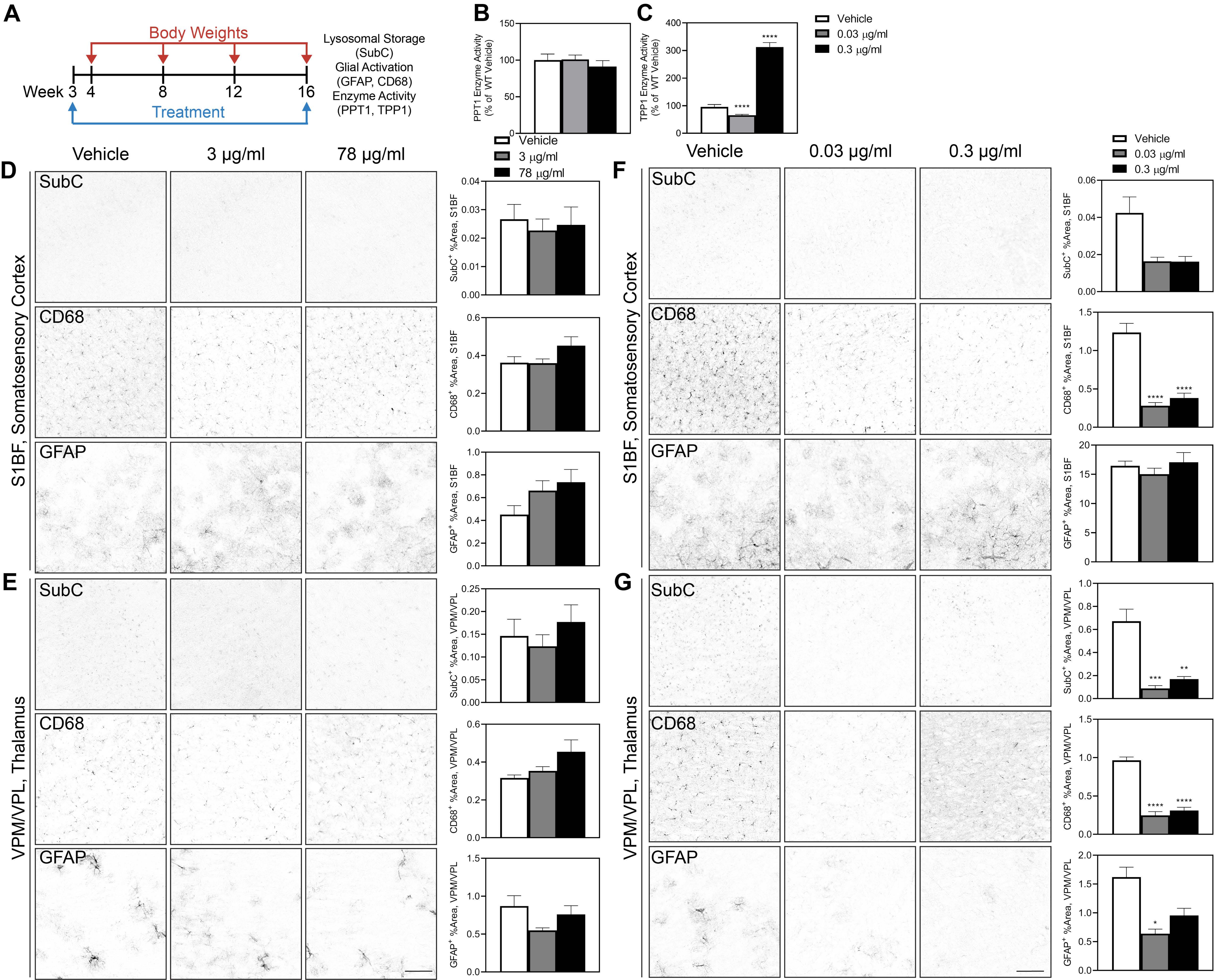
A
Body Weights
Week 3 4 8 12 16
Treatment
Lysosomal Storage (SubC)
Glial Activation (GFAP, CD68)
Enzyme Activity (PPT1, TPP1)
B
PPT1 Enzyme Activity (% of WT Vehicle)
C
TPP1 Enzyme Activity (% of WT Vehicle)
Vehicle
0.03 μg/ml
0.3 μg/ml
Vehicle
3 μg/ml
78 μg/ml
D
S1BF, Somatosensory Cortex
SubC
CD68
GFAP
SubC+ %Area, S1BF
CD68+ %Area, S1BF
GFAP+ %Area, S1BF
E
VPM/VPL, Thalamus
SubC+ %Area, VPM/VPL
CD68+ %Area, VPM/VPL
GFAP+ %Area, VPM/VPL
F
G

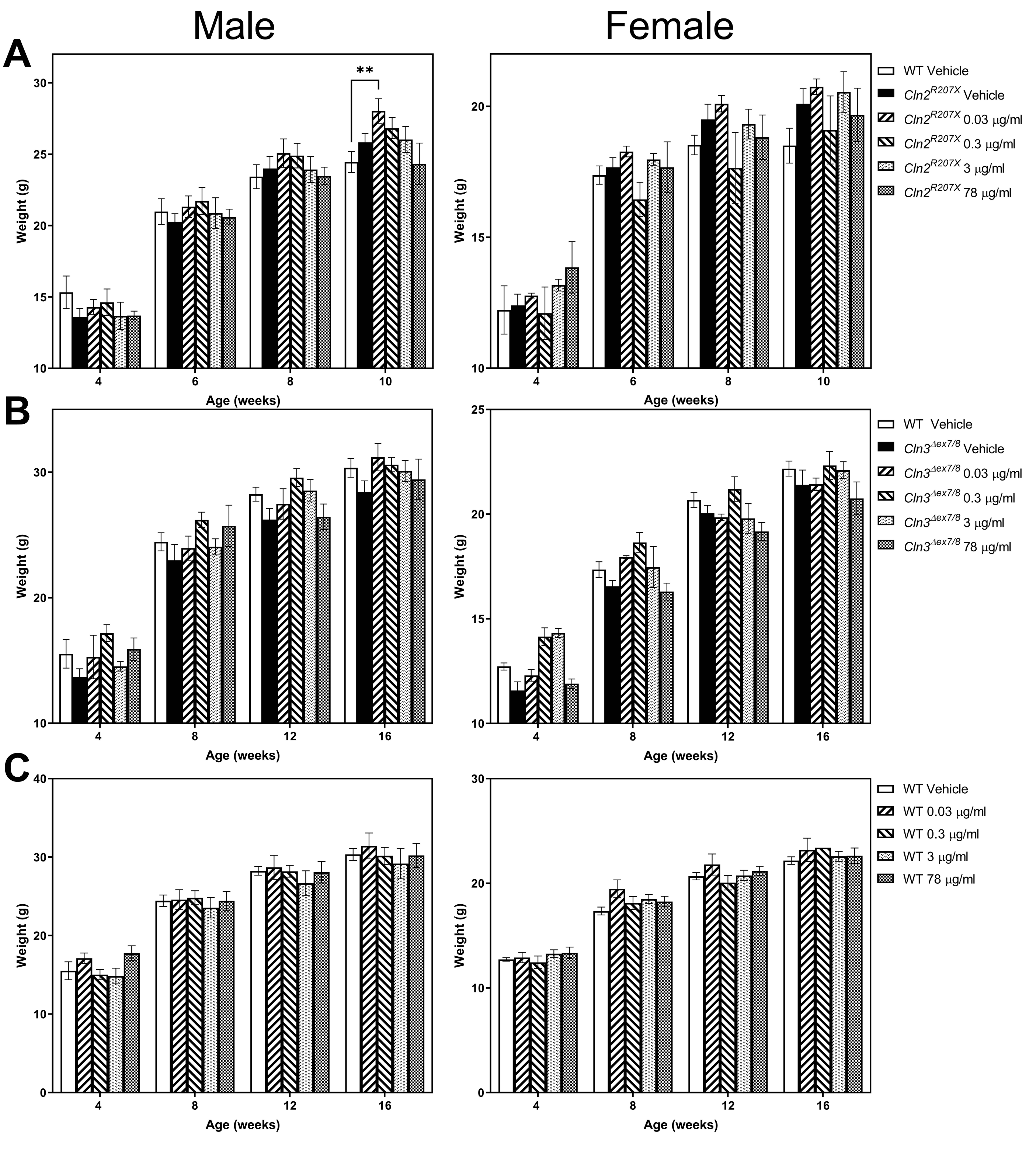

Male
Female
A
B
C
**
Weight (g)
Age (weeks)
WT Vehicle
Cln2R207X Vehicle
Cln2R207X 0.03 μg/ml
Cln2R207X 0.3 μg/ml
Cln2R207X 3 μg/ml
Cln2R207X 78 μg/ml
WT Vehicle
Cln3Δex7/8 Vehicle
Cln3Δex7/8 0.03 μg/ml
Cln3Δex7/8 0.3 μg/ml
Cln3Δex7/8 3 μg/ml
Cln3Δex7/8 78 μg/ml
WT Vehicle
WT 0.03 μg/ml
WT 0.3 μg/ml
WT 3 μg/ml
WT 78 μg/ml
